# OncoGEMINI: Software for Investigating Tumor Variants From Multiple Biopsies With Integrated Cancer Annotations

**DOI:** 10.1101/2020.03.10.979591

**Authors:** Thomas J. Nicholas, Michael J. Cormier, Xiaomeng Huang, Yi Qiao, Gabor T. Marth, Aaron R. Quinlan

## Abstract

DNA sequencing has unveiled extensive tumor heterogeneity in several different cancer types, with many exhibiting diverse subclonal populations. Identifying and tracing mutations throughout the expansion and progression of a tumor represents a significant challenge. Furthermore, prioritizing the subset of such mutations most likely to contribute to tumor evolution or that could serve as potential therapeutic targets represents an ongoing problem. Here we describe **OncoGEMINI**, a new tool designed for exploring the complex patterns and trajectory of somatic and inherited variation observed in heterogeneous tumors biopsied over the course of treatment. This is accomplished by creating a searchable database of variants that includes tumor sampling timepoints and allows for filtering methods that reflect specific changes in variant allele frequencies over time. Additionally, by incorporating existing annotations and resources that facilitate the interpretation of cancer mutations (e.g., CIViC, DGIdb), OncoGEMINI enables rapid searches for, and potential identification of, mutations that may be driving subclonal evolution.

## Introduction

Cancers arise from a variety of genetic alterations and, over time, often accumulate a substantial mutational load leading to subclonal diversity^1,2^. As a result, DNA sequencing of tumors has revealed extensive heterogeneity within primary tumors or between primary and subsequent occurrences^3^, and that the degree of heterogeneity is correlated with patient outcomes^4^. Prioritizing mutations in the face of this heterogeneity relies upon accurate variant discovery and being able to differentiate variants with potential relevance from those that are less likely to contribute to the proliferation of a given tumor. Nevertheless, properly deciphering which identified variants, if any, contribute to tumor origin, survival, and proliferation is crucial to understanding tumor biology and determining potential treatments.

Recognizing this need, we introduce OncoGEMINI as new software to explore genetic variation observed across multiple tumor biopsies and facilitate the identification of both inherited and somatic mutations that may be involved in tumor progression or resistance. OncoGEMINI is uniquely effective in the analysis of variation observed in either a single or multiple cancer biopsies from one or more patients. OncoGEMINI builds upon the GEMINI framework^5^, which creates a database from a VCF^6^ file and extensively annotates genetic variants in an effort to facilitate analysis. GEMINI was designed for the analysis of inherited variants in studies of rare disease^7–9^, and is poorly suited to the analysis of somatic mutations, tumor heterogeneity, and the analysis of multiple biopsies that vary over time and location in the patient’s body.

OncoGEMINI addresses these limitations and enables rapid variant exploration in tumor sequencing studies, especially those featuring longitudinal data across multiple time points from a single patient. To better select or prioritize tumor variants, OncoGEMINI integrates several cancer-relevant genomic annotations that serve as searchable terms to differentiate variants from one another. Additionally, OncoGEMINI provides multiple filtering tools that search for various signatures of tumor heterogeneity, including observable allele frequency changes across multiple samples. These include the `bottleneck`, `loh`, `truncal`, and `unique` tools. By using these filtering tools in combination with specific cancer annotations, OncoGEMINI can effectively prioritize tumor variants that may drive tumor progression or be potential treatment targets.

## Design

### OncoGEMINI framework

OncoGEMINI employs the same general functionality as GEMINI and imports variant information, including sample genotypes, from a VCF file into a searchable SQLite database. OncoGEMINI is intended to be used alongside the VCF annotation tool, vcfanno^10^, and the database creation tool, vcf2db^11^. Together these allow for efficient loading of user specified (or created) annotations to be included in the resulting database (**Figure 1**). Database creation times vary depending on the number of variants and annotations included in the annotated VCF file, but over a thousand variants per second can typically be inserted into the database using vcf2db.

**Figure 1.**
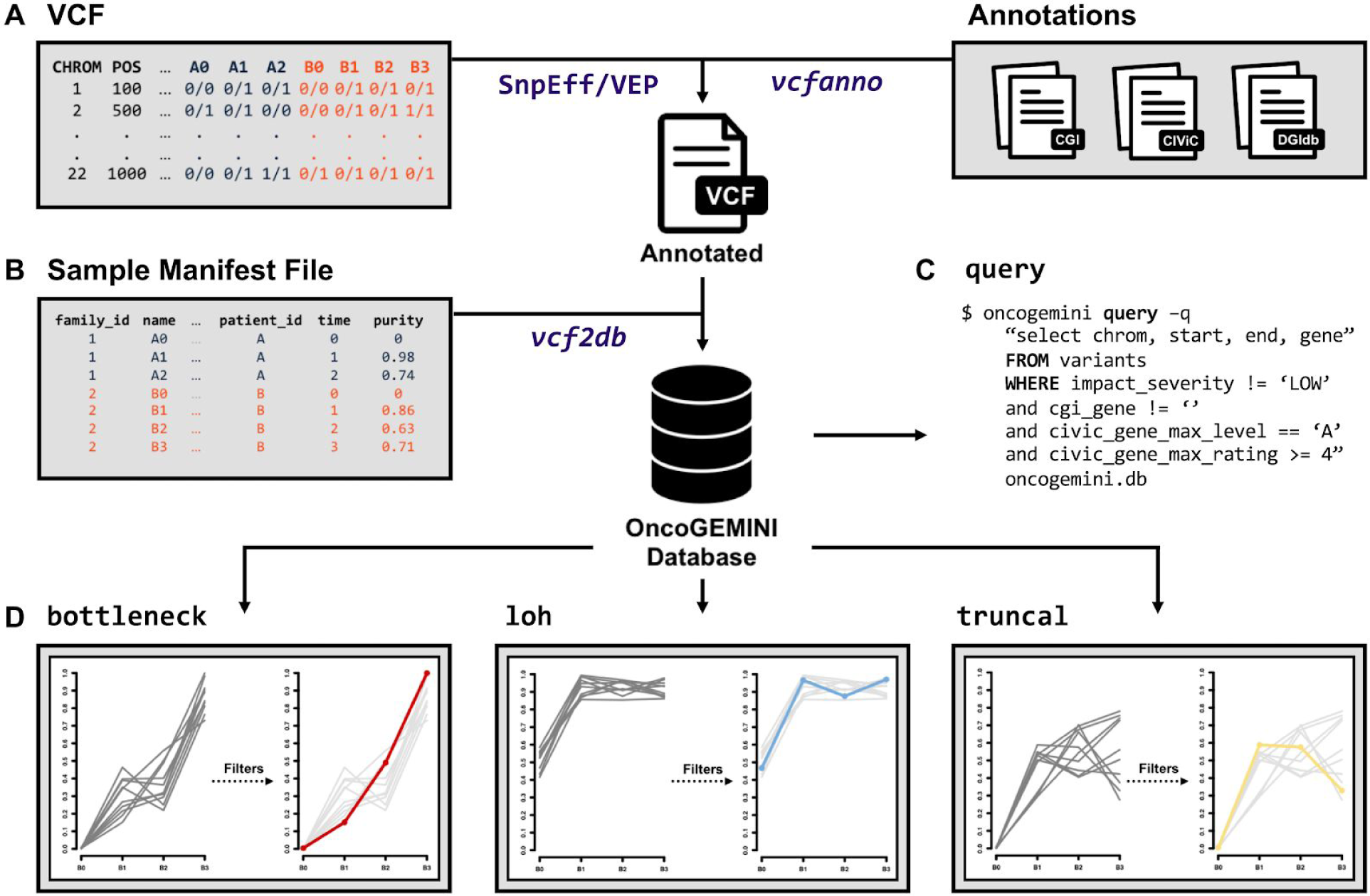
Overview of OncoGEMINI database creation and usage. **A.** Genetic variants from a VCF file are first annotated by the user with either SnpEff or VEP and then additional user-determined annotations can be added to the SnpEff or VEP annotated VCF file using the vcfanno tool. **B.** The fully annotated VCF, along with a sample manifest file, is then used to create an OncoGEMINI database via vcf2db. Once the OncoGEMINI database is created, users are able to **(C)** create their own customized queries with the query command or **(D)** select from built-in tools (such as `bottleneck`, `loh`, or `truncal`) to filter variants. To better select and focus on specific variants users may combine tools and queries with included annotations to further filter the number of variants.

An important change to the existing GEMINI framework that is necessary for some of OncoGEMINI’s functionality is the inclusion of a tumor specific sample manifest file used during database loading. OncoGEMINI utilizes a user generated sample manifest file to describe the distinctive relationships that may exist between tumor samples. This manifest retains the same basic structure of common pedigree files with additional columns describing `patient_id`, `time`, and lastly, `purity`. Given that multiple samples may be derived from the same patient, the `patient_id` column ensures that sample relatedness is properly catalogued. Since these samples may be obtained through a series of biopsies at different timepoints, the `time` column describes the temporal relationship between samples for a given patient. While OncoGEMINI does not compute tumor purity estimates, if such estimates are known they may be included in the `purity` column and invoked as a command line parameter to alter allele frequencies accordingly. Altogether, these extra columns give OncoGEMINI greater flexibility and specificity when performing queries or employing different filtering schemes.

### Cancer-specific annotations

Diverse databases and annotations have been developed to help interpret identified tumor variants with their specific relevance to various aspects of tumor origin, growth, and treatment^12–16^. Such annotations can be essential in determining which identified tumor variants are actionable or otherwise merit further investigation from those that are less relevant to tumor progression. Many of these annotation sources exist independently of one another and not always in formats that lend themselves readily to bioinformatic applications.

We have integrated a number of cancer relevant annotation resources including the Cancer Genome Interpreter (CGI)^17^, Clinical Interpretations of Variants in Cancer (CIViC)^18^, and the Drug Gene Interaction database (DGIdb)^19^. Each annotation provides pertinent information relating to various aspects of tumor biology and, in some cases, have also summarized various genome annotations from other resources. We have made all of these annotation files, and guides to their creation, available via the Cancer Relevant Annotations Bundle, CRAB, resource (https://github.com/fakedrtom/crab). While the relevance of each annotation varies, especially with regards to particular cancer types, they provide valuable insights into the pathogenicity and frequency of genetic variants, the propensity for certain genes to harbor mutations in specific cancer types, or known drug susceptibilities and interactions with genes or individual variants. From each of these we have converted selected information into a series of summarized files that can then be used to annotate a VCF file in preparation for loading into a OncoGEMINI database. Given the inherent flexibility of vcfanno, any combination of these annotations or any additional ones not included here, may be selected and added to a VCF. We expect that this will be a valuable resource even outside of use within OncoGEMINI and are eager to add more annotations, references, and data to improve its utility.

### OncoGEMINI functionality

Combining tumor variants with genome annotations into a OncoGEMINI database enables the identification and prioritization of tumor relevant genetic variants. Once loaded, the database is populated with the information contained within the selected VCF. Variants from the VCF become rows in the database and annotations are stored as columns within the OncoGEMINI database that can be useful for isolating variants of interest.

### The *query* tool

Via OncoGEMINI’s `query` tool, users are able to impose specific search parameters to identify variants meeting custom search criteria. For example, if one wanted to filter all variants in a OncoGEMINI database to only those with the highest validated clinical association as documented in CIViC (i.e., an evidence level of ‘A’), the following query would restrict the output to the genomic coordinates, reference and alternate alleles, and gene (if applicable), for all variants that match those in the CIViC database with an evidence level of ‘A’:

~~~
oncogemini query -q “select chrom, start, end, ref, alt, gene from variants
where civic_evi_level = ‘A’” oncogemini.db
~~~

The `query` tool can take advantage of the flexibility of OncoGEMINI’s SQL language to build sophisticated search commands. Anticipating common searches pertaining to tumor growth and propagation, we have also developed a suite of tools that identify variants that follow patterns of particular interest in tumor evolution and alleviate the need to repeatedly craft lengthy search queries. Each of the tools described in the following sections are capable of further customization with additional tool specific parameters and via the annotations included in the database. Together, these options provide the ability to design explicit search queries that can effectively reduce the list of variants, thereby narrowing in on potential variants of interest.

### The *bottleneck* tool

As individual tumor cells replicate, they may acquire private mutations that give rise to subclonal structures and heterogeneity within tumors. Over time, owing to drift or the selective pressure of drug and other treatments, the prevalence of certain tumor variants can be reduced or increased. Thus, the ability to determine the frequency at which any given tumor variant exists is both a function of sampling time and efficiency. Variants which increase in frequency over time, especially across different interventions, may represent potential contributors to the proliferation and survivability of a given tumor. We have developed the `bottleneck` tool to identify variants exhibiting this increased allele frequency over time. This tool relies upon having information regarding the sequential order that tumor samples arose; therefore, this tool is most applicable to longitudinal data, but may be useful with other studies, including metastatic studies, provided temporal information is otherwise appropriately known or ascertained. The `bottleneck` tool scans each variant in the database for samples that exhibit increasing allele frequencies over the sampling time indicated in the sample manifest file (**Figure 2**). To accomplish this, the `bottleneck` tool uses the allele frequencies of specified samples across the sample timepoints and calculates the slope of these values. Variants that exceed a given slope (default `0.05`) are reported in the output.

**Figure 2.**
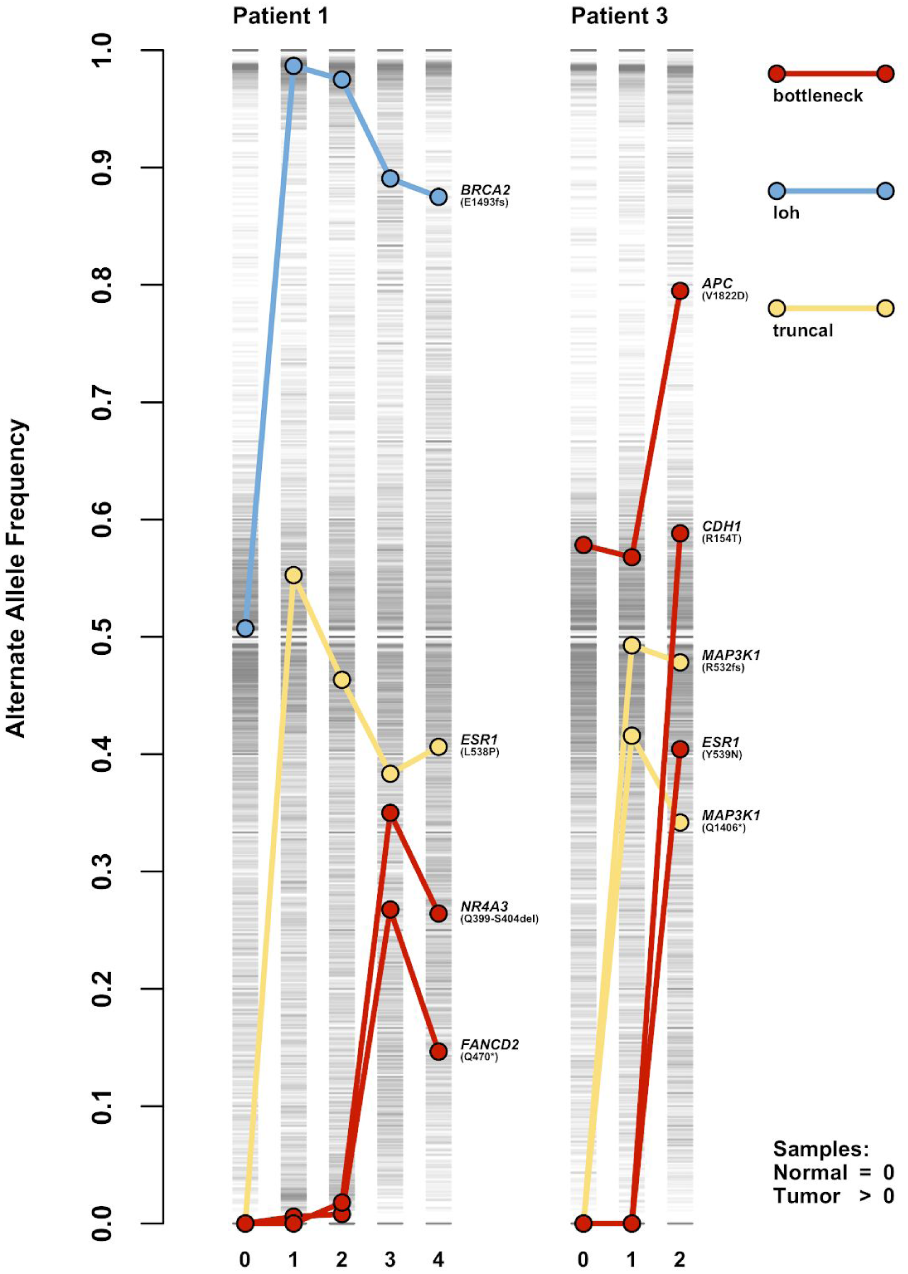
Using OncoGEMINI filtering tools to identify variants with specific patterns. Genetic variants corresponding to all samples originating from representative patients, #1 and #3 (further details in “Using OncoGEMINI” section), are indicated with single gray lines at their respective alternate allele frequencies, where darker grays represent a greater accumulation of variants with similar frequencies. Samples are represented by their sampling time as indicated in the sample manifest file, where a time of `0` indicates a normal tissue sample and values greater than `0` correspond to subsequent tumor samples. Individual variants and their alternate allele frequency changes across multiple tumor timepoints are highlighted with colored points connected by lines and were isolated from all other variants using the OncoGEMINI filtering tools: `bottleneck` (red), `loh` (blue), and `truncal` (yellow). For each of these variants, the gene in which they occur and their specific mutation are listed next to the point in the final sample for each patient.

### The *loh* tool

Loss of heterozygosity (LOH) is a common genomic alteration in cancer genomes that leads to the loss of one allele. LOH events can result in functional consequences including reduced gene expression, haploinsufficiency, or acting as the second “hit” in a tumor suppressor gene. LOH mutations can indicate causative variants or otherwise serve as biomarkers for cancer identification and potential patient care. One method for identifying LOH relies upon observing genetic loci that appear heterozygous in germline DNA but are not heterozygous in tumor DNA. The `loh` tool identifies potential LOH variants by observing allele frequency changes that are consistent with heterozygosity present in the normal tissue samples, but absent in all specified tumor samples (**Figure 2**).

### The *truncal* tool

Somatic mutations that arose early in tumor development are likely to be present in all sequenced samples. Such mutations may serve as important therapeutic targets since they are present throughout all samples, rather than localized and private to any specific subclone(s). For example, this subset of variants may harbor potential neoantigens which represent ideal targets for patient-specific T cell-based cancer immunotherapy^20^. The `truncal` tool was designed to identify genetic variants that are present in all given tumor samples, but absent from any normal samples (**Figure 2**). Similar to the `loh` tool, the `truncal` tool also requires a normal sample to be included and accepts a defined maximum allele frequency to be allowed in the normal tissue samples (default is `0`). Variants where all the tumor samples have allele frequencies that are higher than the maximum normal tissue allele frequency are included in the output.

### The *unique* tool

While one of the key features of analyses with OncoGEMINI is the inclusion of multiple samples from different spatial and temporal biopsies, the `unique` tool also allows OncoGEMINI to highlight variants that are specific to a single sample or group of samples. This tool requires a desired sample(s) be listed and returns all variants that are present in that sample(s) and absent from all others. Similar to the `truncal` tool, this is done by specifying a minimum allele frequency (default is `0`) that must be exceeded in the listed sample(s), but not be met in all other samples.

### Identifying somatic mutations

OncoGEMINI will evaluate all variants within the database and select those that meet specified tool and annotation filter requirements. Thus, if the VCF used to create the database contained both germline and somatic mutations, both mutation types would be considered by OncoGEMINI commands. To focus solely on somatic mutations, it is recommended that the VCF used for the creation of a OncoGEMINI database be pre-filtered to only include somatic mutations or that somatic mutations be clearly labeled in the VCF so they are incorporated as a filterable annotation within the database. If that is not possible, the `set-somatic` tool may be employed which allows for variants within a OncoGEMINI database to be “flagged” as somatic based on user defined criteria regarding normal and tumor sample sequencing depths and allele frequencies. OncoGEMINI tools may then take advantage of the `--somatic-only` parameter to restrict variant evaluations to only those variants that have been marked as somatic in the database by the `set-somatic` tool.

## Using OncoGEMINI

OncoGEMINI’s integration of genetic variants identified across one or more biopsies with annotations relevant to cancer enables a wide range of analyses and variant prioritization strategies. Furthermore, the OncoGEMINI framework can be used to study variation observed in diverse study designs, including: studies of a single tumor and normal biopsy and multiple tumor and/or ascites biopsies from the same patient over the course of treatment. Existing tools^21–25^ are well-suited to the study of mutations found in matched tumor-normal studies and OncoGEMINI’s primary innovation is the ability to analyze multiple related samples alongside one another. We therefore demonstrate OncoGEMINI’s functionality by providing an example analysis that isolates variants of interest in a recent study of longitudinal biopsies from three breast cancer patients^26^.

### Breast Cancer Longitudinal Data

We specifically focused on patients #1 and #3, each of which were followed for more than 3 years and experienced multiple tumor recurrences (4 total tumor samples in patient #1 and 2 in patient #3), undergoing biopsies with samples taken before, during, or after a variety of distinct drug treatments. Specific details regarding these patients, their tumor history, treatments, data generated and conducted analyses were previously reported^26^, but in summary, the original study reported mutations that were identified through a combination of manual curation, analysis, and validation; automating future analyses as much as possible was a key motivation for the development of OncoGEMINI. Previously, nine total SNV or small indel variants were identified in patients #1 and #3 and these mutations serve as ideal candidates for the types of variants of interest that OncoGEMINI should be able to prioritize using its improved annotations and filtering tools (for specific variant details, see **Figure 2**).

For each VCF, variants were annotated in two distinct steps. First, consistent with GEMINI, OncoGEMINI also incorporates standard variant effect predictions made by SnpEff^27^ or VEP^28^, and in this case, initial annotations were added via SnpEff. Second, we downloaded additional cancer-relevant features from various databases, including CGI, CIViC, and DGIdb, via the CRAB and subsequently added them as variant annotations to the VCFs using vcfanno. A sample manifest file was prepared that listed all of the samples, their names, the patients they corresponded to (either patient #1 or #3) and timepoints where the normal samples were given a time of `0` and subsequent tumor samples were given times greater than `0` with each consecutive sample time increasing by `1`. The optional `purity` column was not included for either of these patients. With the sample manifest file, OncoGEMINI databases, including all of the added annotations, were then created for each patient’s VCF using vcf2db. Resulting databases contained 5,543,181 and 5,355,078 total variants for patients #1 and #3, respectively, which are reduced to 4,986,519 and 4,928,034 if we require a minimum sequencing depth threshold of 10 for all samples.

To further reduce this number and reveal potential variants of interest (i.e., the previously reported variants) we required certain filters via the included annotations to be met and utilized many of the previously described OncoGEMINI tools. As expected, the count of variants that OncoGEMINI returns depends upon the number of annotation filters and tools that are specified where, generally speaking, the more filters alongside tools that are required, the more the list of returned variants is reduced.

Specifying that variants be filtered to only those with a SnpEff impact prediction of “medium” or “high” (`impact_severity != ‘LOW’`), drastically reduced the number of returned variants to 20,580 and 19,516. Also restricting variants to those found in genes with previous implications towards tumorigenesis in certain cancer types via CGI’s Catalog of Cancer Genes (`cgi_gene != ‘’`) and genes with high CIViC evidence levels or ratings (`civic_gene_max_level == ‘A’ or civic_gene_max_level == ‘B’ or civic_gene_max_rating >= 4`) refined the number of variants to 140 and 149. Combining these annotation filters with each of the OncoGEMINI `bottleneck`, `loh`, and `truncal` tools resulted in a total of three variants from patient #1 and 2 variants from patient #3 being returned which accounted for three of the nine previously reported variants. To try and recover the remaining 6 previously reported variants, we relaxed the filter requirements by removing the CIViC evidence level and rating annotation filters. This increased the number of resulting variants to 1,245 and 1,253, respectively, but by once again passing them through the `bottleneck`, `loh`, and `truncal` tools only 17 and 27 variants remained, including eight of the nine previously reported variants. Altogether with just a few commands that returned results in seconds of time, we were able to filter millions of tumor variants to a much more manageable number in each patient, which included nearly all of the previously reported variants (**Figure 3**). We note that distinct annotations from SnpEff, CGI, and CIViC are each capable of substantially reducing the number of variants, but by combining these annotations filters with one another and each of the OncoGEMINI filtering tools the number of variants is refined to a specific few. We also identified in each patient, additional mutations that were not previously reported, but that meet the same parameters as those that were. While these variants are not validated and their potential role in the subclonal tumorigenesis in each of the patients is unknown, they may be worth further consideration.

**Figure 3.**
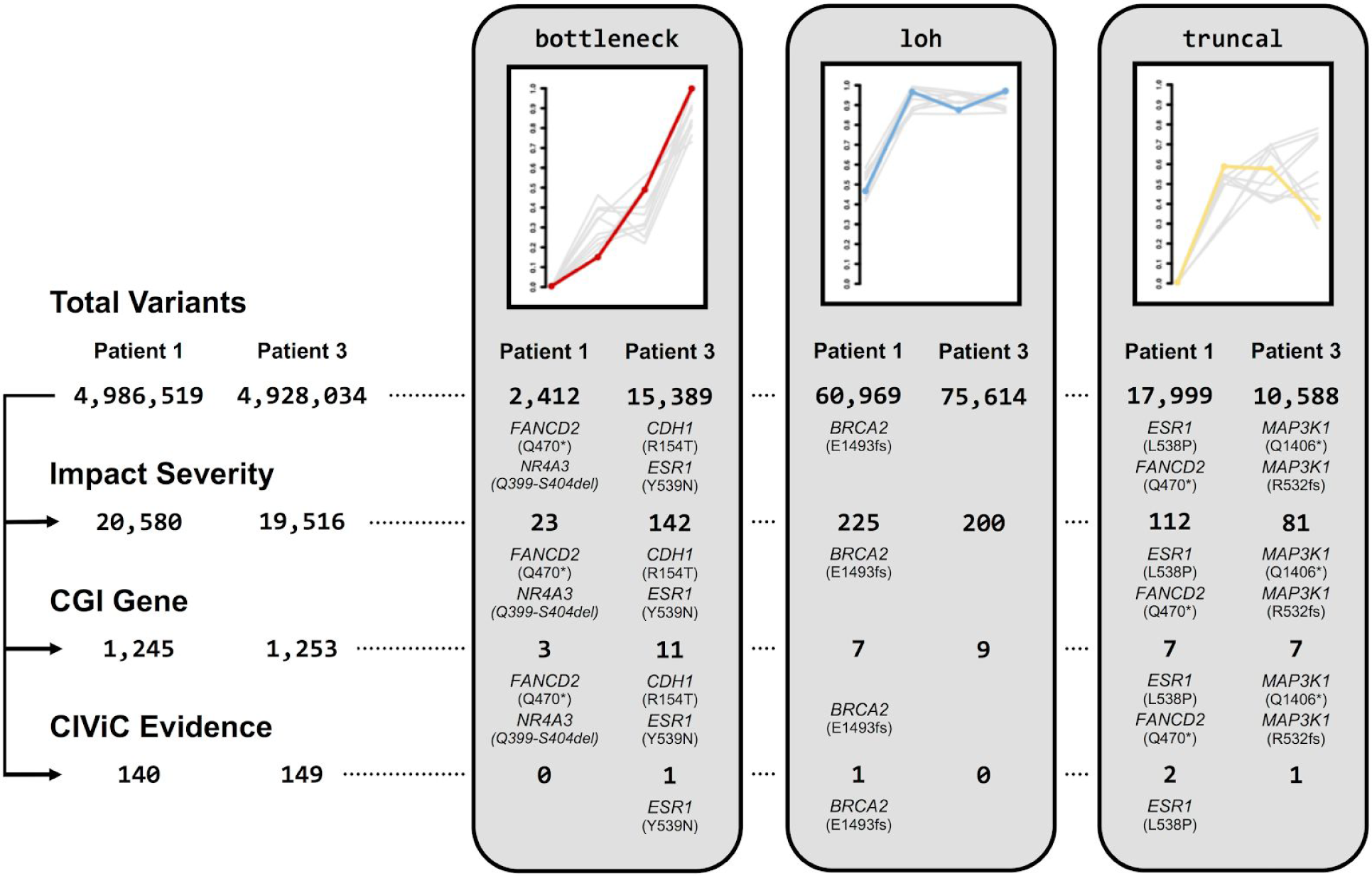
Schematic of an OncoGEMINI workflow for filtering variants. Variants are filtered using a combination of OncoGEMINI tools and included annotations as filter requirements. For each set of numbers, those listed on the left belong to patient #1 and those on the right are from patient #3. On the leftmost side of the figure, the total number of variants for each patient are listed with an increasing number of cancer annotation filter requirements being added from top to bottom, and the corresponding number of variants that meet those requirements, being specified as individual rows. For each row, the number of variants is further restricted by individual OncoGEMINI tools listed towards the right side of the figure, highlighted in light gray boxes. The number of variants returned by different combinations of OncoGEMINI tools and cancer annotation filters are continued as rows within each of the gray boxes corresponding to different tool types. Any combinations of OncoGEMINI tools and cancer annotation filters that returned any of the previously identified variants are specified with the genes in which the variants were found and their specific mutations.

It is important to note that while this filtering scheme has been described in a sequential manner, all of the searches, including the application of multiple filters with individual tools, can be accomplished with a single command. For example, using the `truncal` tool alongside all of the previously described filters can be done like so:

~~~
$ oncogemini truncal --minDP 10 --columns “chrom,start,end,ref,alt,gene”
--filter “impact_severity != ‘LOW’ and cgi_gene != ‘’ and
(civic_gene_max_level == ‘A’ or civic_gene_max_level == ‘B’ or
civic_gene_max_rating >= 4)” patient1.db
~~~

This command will return the chromosome, start and end positions, reference and alternate alleles, and the gene for the two truncal variants in patient #1 that meet these criteria. This allows for maximum customization into a single command when devising search specifications.

## Conclusions and Discussion

Building upon the GEMINI framework, OncoGEMINI is ideally suited to the exploration and prioritization of tumor mutations. Cancer research increasingly requires the integration of data from not only multiple biological samples, but also the accumulated genomic information that is found in a variety of distinct databases. We have designed OncoGEMINI to combine information from numerous sources alongside longitudinal tumor sequence data to enable the rapid identification of variants that match specific patterns and criteria. Therefore, OncoGEMINI provides a unique tool that assists in complex cancer analyses.

While much of OncoGEMINI’s functionality is applicable to a variety of data from different tumor studies, it is optimized to incorporate and analyze data from multiple tumor samples belonging to the same patient and is thus most powerful when used with longitudinal data that has been collected over various time points and treatments. OncoGEMINI offers a number of specific filtering tools that focus on variant allele frequency changes between samples and each tool can be further adjusted with included options and parameters. Additionally, using the vcfanno tool, user defined OncoGEMINI databases can be built that enable different genomic annotations to be incorporated and used in identifying relevant tumor mutations, thus empowering powerful cancer-specific queries. We have suggested and prepared specific cancer annotations, but given the inherent flexibility afforded by vcfanno, custom annotations provided or created by users can be included. This allows OncoGEMINI greater versatility and suitability to a wide range of cancer analysis projects. By combining annotation information with OncoGEMINI tools, we demonstrated that individual tumor variants can be identified with simple commands that run quickly in a matter of seconds.

Even though OncoGEMINI is not clinically deterministic, it can help rapidly sort through tumor variants from sequencing data and identify individual variants fitting specific requirements that may be indicative of variants with potential clinical significance. The default settings of the OncoGEMINI tools aim to pinpoint such likely relevant variants, but being primarily an exploratory tool, OncoGEMINI’s individual tool settings can be adjusted, as is appropriate, to expand query criteria. For example, the last remaining variant from the previously discussed longitudinal example was not recovered in the described analysis (patient #3, *APC* V1822D) because it was a germline variant that increased in frequency over the time course of the tumor samples (**Figure 2**). This variant follows the allele frequency pattern that is similar to what is expected to be found by the `bottleneck` tool, however, the default settings of the `bottleneck` tool would ignore this variant because of its high allele frequency in the included germline (normal) tissue. By adjusting the defaults and allowing for a larger initial normal tissue allele frequency using the `--maxNorm` parameter (or by omitting the normal sample altogether from the analysis using the `--samples` parameter), we are able to recover that previously reported variant as well, but further filtering would be required to narrow in on this specific mutation. In this manner, OncoGEMINI enjoys further flexibility that enhances its capabilities as a tool for surveying tumor heterogeneity from sequencing data.

OncoGEMINI is an open-source software package and it is freely available. Source code and further documentation can be found at: https://github.com/fakedrtom/oncogemini.

